# SLICER: Inferring Branched, Nonlinear Cellular Trajectories from Single Cell RNA-seq Data

**DOI:** 10.1101/047845

**Authors:** Joshua D. Welch, Alexander Hartemink, Jan F. Prins

**Affiliations:** Department of Computer Science, University of North Carolina at Chapel Hill; Curriculum in Bioinformatics and Computational Biology, University of North Carolina at Chapel Hill; Department of Computer Science, Duke University; Program in Computational Biology and Bioinformatics, Duke University

**Keywords:** Single cell RNA-seq, time series, manifold learning

## Abstract

Single cell experiments provide an unprecedented opportunity to reconstruct a sequence of changes in a biological process from individual “snapshots” of cells. However, nonlinear gene expression changes, genes unrelated to the process, and the possibility of branching trajectories make this a challenging problem. We developed SLICER (Selective Locally Linear Inference of Cellular Expression Relationships) to address these challenges. SLICER can infer highly nonlinear trajectories, select genes without prior knowledge of the process, and automatically determine the location and number of branches and loops. SLICER more accurately recovers the ordering of points along simulated trajectories than existing methods. We demonstrate the effectiveness of SLICER on previously published data from mouse lung cells and neural stem cells.

## Introduction

Understanding the dynamic regulation of gene expression in cells requires the study of important temporal processes, such as cell differentiation, the cell division cycle, or tumorigenesis. However, in such cases, the precise sequence of changes is generally not known, few if any marker genes are known, and individual cells may proceed through the process at different rates. These factors make it very difficult to externally judge where a cell is in the process. Additionally, bulk RNA-seq data may blur aspects of the process because cells at sampled at a given point in time may be at different points in the process.

The advent of single cell RNA-seq enables the study of sequential gene expression changes by providing a set of time slices or “snapshots” from individual cells sampling different moments in the process [1‒3]. To combine these snapshots into a coherent picture, we need an “internal clock” that tells, for each cell, where it is in the process. Because one of the motivations for performing a single cell RNA-seq experiment is to conduct an unbiased, genomewide study, we would like an unsupervised approach for inferring this internal clock, rather than relying on known marker genes or experiments starting from synchronized cells. Given these motivations, the internal state of a cell is the only reliable way to judge where it is in the process.

One way to approach this problem is to infer a low-dimensional manifold embedded in high-dimensional space that captures the observed geometric relationships among the cells [1, 2]. The modeling assumption behind this approach is that the main difference among cells is where they lie in the process, so that the sequence of gene expression changes traverses a “trajectory” through the sampled cells in high-dimensional space.

Several techniques to identify cellular trajectories have recently been developed. The Monocle tool [1] uses independent component analysis (ICA) to find a low-dimensional linear projection of the data, then constructs a minimum spanning tree in the resulting low-dimensional space to order cells progressing through development. Another tool, Wanderlust, constructs an ensemble of *k*-nearest neighbor graphs directly in high-dimensional space without performing dimensionality reduction, then finds shortest paths through the ensemble of graphs [2]. An advantage of Wanderlust is its ability to capture nonlinear behavior.

Monocle and Wanderlust have both been successfully applied to reveal biological insights about cells moving through a biological process [1, 2, 4, 5]. However, a number of aspects of the trajectory construction problem remain unexplored. For example, both Monocle and Wanderlust assume that the set of expression values they receive as input have been curated in some way using biological prior knowledge. Wanderlust was designed to work on data from protein marker expression, a situation in which the number of markers is relatively small (dozens, not hundreds of markers) and the markers are hand-picked based on prior knowledge of their involvement in the process. In the initial application of Monocle, genes were selected based on differential expression analysis of bulk RNA-seq data collected at initial and final timepoints [1]. In addition, Monocle uses ICA, which assumes that the trajectory lies along linear projection of the data. In biological settings, this assumption may not hold. In contrast, Wanderlust can capture nonlinear trajectories, but works in the original high-dimensional space, which may make it more susceptible to noise, particularly when given thousands of genes, many of which are unrelated to the process being studied. Another challenging aspect of trajectory construction is the detection of branches. For example, a developmental process may give rise to multiple cell fates, leading to a bifurcation in the manifold describing the process. Wanderlust assumes that the process is nonbranching when constructing a trajectory. Monocle provides the capability of dividing a trajectory into a branches, but requires the user to specify the number of branches.

In this paper, we present SLICER (Selective Locally linear Inference of Cellular Expression Relationships), a new approach that uses locally linear embedding (LLE) to reconstruct cellular trajectories. SLICER provides four significant advantages over existing methods for inferring cellular trajectories: (1) the ability to automatically select genes to use in building a cellular trajectory with no need for biological prior knowledge; (2) use of locally linear embedding, a nonlinear dimensionality reduction algorithm, for capturing highly nonlinear relationships between gene expression levels and progression through a process; (3) automatic detection of the number and location of branches in a cellular trajectory using a novel metric called geodesic entropy; and (4) the capability to detect types of features in a trajectory such as "bubbles" that no existing method can detect.

## Results

### Overview of SLICER Method

Figure 1 summarizes the process by which SLICER infers cellular trajectories. SLICER takes as input a matrix of unfiltered gene expression levels. By computing a quantity we term “neighborhood variance”, we choose a set of genes to use in building the trajectory (Fig. 1a). Intuitively, this method removes genes that show random fluctuation across the set of cells and selects only genes that vary incrementally from cell to cell in a systematic manner. Note that this gene selection method does not require either prior knowledge of genes involved in the process or differential expression analysis of cells from multiple time points. Next, the number of nearest neighbors *k* to use in constructing a lowdimensional embedding is chosen so as to yield the shape that most resembles a trajectory, as measured by the *α*-convex hull of the embedding (Fig. 1a and Fig. S1). Alternatively, the user can specify *k* to manually tune the trajectory. SLICER then uses a nonlinear dimensionality reduction algorithm, locally linear embedding (LLE), to project the set of cells into a lower-dimensional space (Fig. 1b). The low-dimensional embedding is used to build another neighbor graph, and cells are ordered based on their shortest path distances from a user-specified starting cell. SLICER then computes a metric called geodesic entropy based on the collection of shortest paths from the starting cell and uses the geodesic entropy values to detect the presence, number, and location of branches in the cellular trajectory (Fig. 1c and Fig. S2). The branch detection approach is based on the insight that the shortest paths along a non-branching trajectory will be highly degenerate, passing through only a small set of cells, in contrast with a branching trajectory which will use one or more distinct sets of cells (see Methods for details).

**Figure 1.**
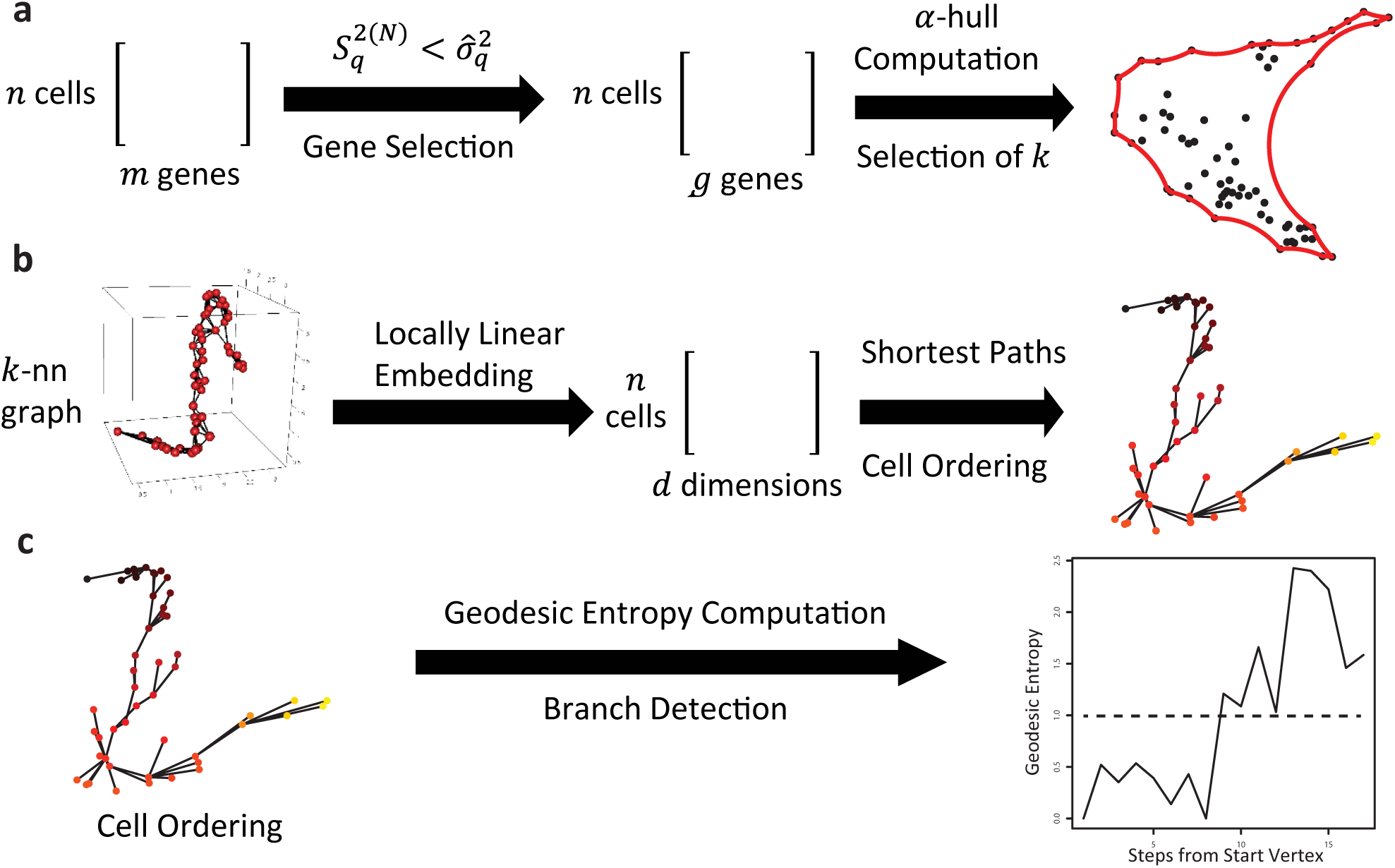
Overview of SLICER method. (a) Genes to use in building a trajectory are selected by comparing sample variance and neighborhood variance. Note that this gene selection method does not require either prior knowledge of genes involved in the process or differential expression analysis of cells from multiple time points. Next, the number of nearest neighbors *k* to use in constructing a lowdimensional embedding is chosen so as to yield the shape that most resembles a trajectory, as measured by the a-convex hull of the cells. (b) SLICER builds a *k*-nearest neighbor graph in highdimensional space and then performs LLE to give a nonlinear low-dimensional embedding of the cells. The low-dimensional embedding is then used to build another neighbor graph, and cells are ordered based on their shortest path distances from a user-specified starting cell. (c) SLICER computes geodesic entropy based on the collection of shortest paths from the starting cell and uses the geodesic entropy values to detect branches in the cellular trajectory.

### Synthetic Data

We constructed a set of simulated trajectories to assess the performance of SLICER on inputs with known solutions. To do this, we generated simulated expression levels for genes in such a way that the expression levels are a function of a “process time” parameter *t*. We simulated 5 different “pathways” using distinct families of functions; the genes generated by a single family of functions are analogous to co-regulated genes in a biological pathway that all change in response to a common regulatory mechanism. Because each simulated gene depends on *t*, points simulated in this way lie along an essentially one-dimensional manifold (a trajectory) in high-dimensional space. Since, in the real data setting, we do not know in advance which genes are involved in a trajectory, we also devised a means to simulate genes that are unrelated to the process. To do this, we randomly permute the simulated values of some genes, thus removing their relationship with *t*. The number of such randomly reshuffled genes is controlled by a parameter *p*.

To measure the performance of a trajectory reconstruction algorithm, we use the algorithm to produce an ordering of the points, then compare it to the true ordering specified by parameter *t*. We used “percent sortedness”, the percentage of pairs of items out of order in a list, as a metric for assessing trajectory reconstruction.

Using the synthetic data generated in this way, we compared SLICER to Wanderlust, a previously published method that can reconstruct nonlinear trajectories. Wanderlust requires the user to specify a value for *k,* the number of nearest neighbors; to ensure a fair comparison, we ran Wanderlust for all values of *k* in [5,10,…,45,50] and chose the *k* that gave the best value. We evaluated SLICER in the same way (testing a sequence of *k* values) and compared the best *k* to the *k* that SLICER automatically selected using our *β*-convex hull approach. To test the importance of using a nonlinear method, we also used ICA, a method that finds a linear projection, to perform dimensionality reduction, then performed the same shortest path algorithm that SLICER uses to order the points in the resulting low-dimensional space. For a baseline method, we randomly permuted the elements in the trajectory and measured the sortedness of the result.

Figure 2 shows the results of this comparison. Several things are important to note about these results. First, Wanderlust performs well when the majority of genes are related to the trajectory, but performance begins to degrade as more unrelated genes are added. This performance degradation may stem from the fact that Wanderlust operates in the original, high-dimensional space, so a large number of irrelevant features begin to compromise the result. In contrast, both SLICER and ICA are fairly stable in the presence of irrelevant genes. However, the ability to capture nonlinear behavior appears to be important, as the performance of ICA is far worse than SLICER and Wanderlust (though still better than a random strategy). Finally, the *α*-hull approach for automatic selection of *k* appears to work well.

**Figure 2.**
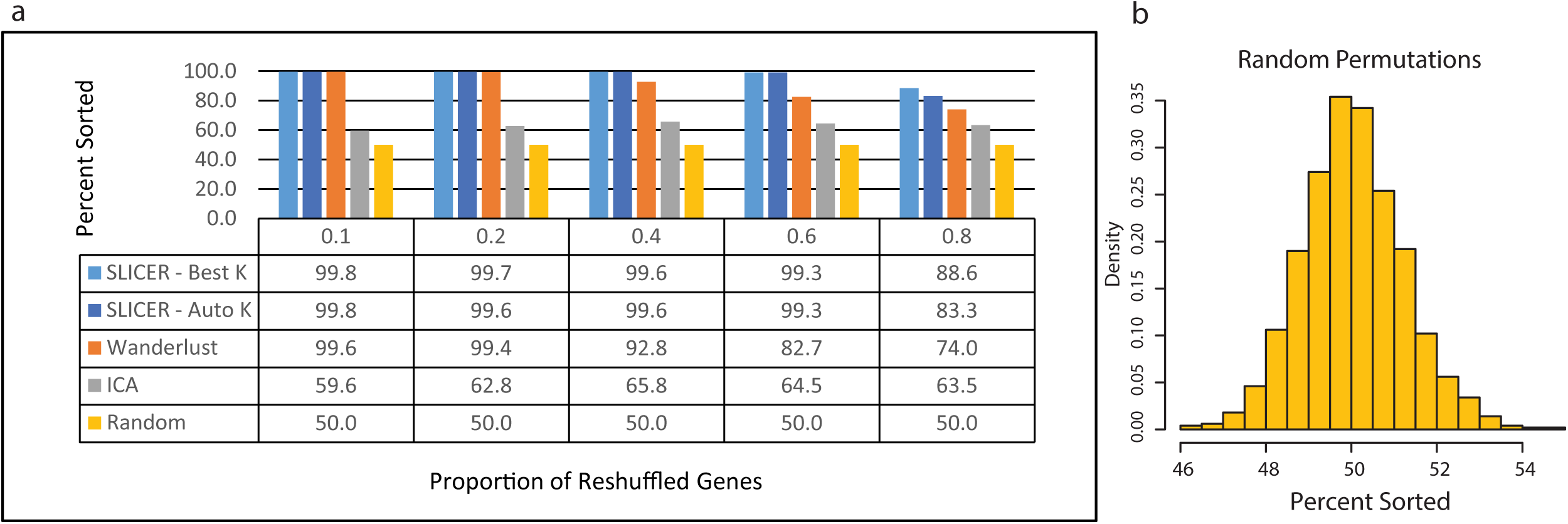
Evaluation of SLICER on synthetic data. (a) Comparison of performance of SLICER, Wanderlust, ICA, and random shuffling. The synthetic datasets were generated as described in the text using 500 genes, *σ* = 2 (*σ* is the noise level), and increasing values of *p*. A higher *p* corresponds to an increased probability that a gene will be randomly reshuffled, removing its relationship with the simulated trajectory. To assess the effectiveness of automatic determination of *k*, SLICER was run both with and without automatic selection of *k*. Performance was evaluated by counting the number of inversions in the resulting sorted list of cells. (b) Histogram of percent sortedness values from 1000 random permutations of the simulated trajectory used in panel a. Note that the distribution of values is sharply peaked around 50% sortedness.

The large performance gap between SLICER and the other methods in Fig. 2 is due in part to the highly curved shape of the trajectory and the use of gene selection. ICA performs poorly on this example because of the large departure from linearity, and both Wanderlust and ICA suffer from the noise added by irrelevant genes. We note, however, that the ability to automatically select relevant genes and reconstruct highly nonlinear trajectories are key benefits of SLICER compared to existing methods. When we simulated a less highly curved trajectory and fed the genes selected by SLICER to the other methods (Fig. S3), the gap between methods was much smaller. SLICER with gene selection and Wanderlust with SLICER’s selected genes were very similar as the proportion of irrelevant genes increased, although Wanderlust performed slightly better in some cases (Fig. S3c). Both methods generally performed better than ICA, with the gap widening as the proportion of irrelevant genes increased (Fig. S3c). SLICER with no gene selection consistently outperformed the other approaches without gene selection (Fig. S3c), highlighting the robustness that LLE provides. We also compared SLICER with the other methods for increasing levels of noise with *p* = 0, that is, no irrelevant genes (Fig. S3d). This comparison showed that the performance of SLICER degrades slightly less rapidly than the other methods in the presence of increasing noise (Fig. S3d), once again indicating the robustness of LLE.

To demonstrate SLICER's ability to detect branches and “bubbles”, we simulated a trajectory in which a single initial path splits into two branches that subsequently converge to a single path (Fig. 3a). We also created a simulated trajectory with a single branch (Fig S4). We created three families of genes in a manner similar to what is described in the Methods section and used 300 genes, noise level *σ* = 0.5, and *p* = 0. Note that the effect of the noise level depends on the relative magnitudes of the genes and the mean of the normal distribution used to add noise. The functions used to simulate the bubble example (Fig. 3) have a much smaller range than the genes used in Fig. 2, and thus a noise level of 0.5 represents a significant challenge (note the level of noise present in Fig. 3a). Fig. 3a contains an example of the three different gene “shapes” used in the simulated dataset.

**Figure 3.**
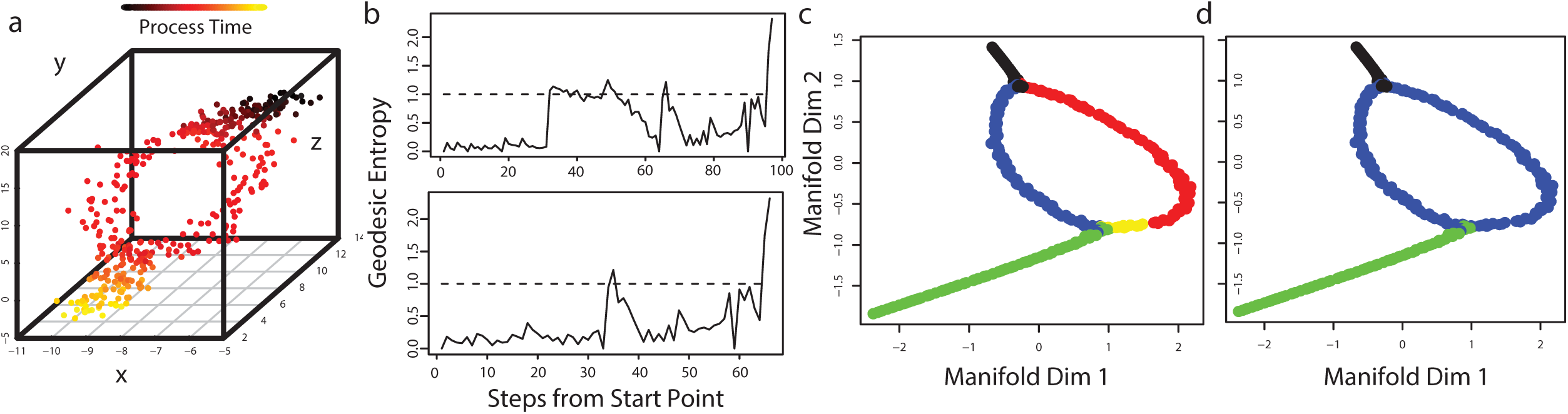
Synthetic data example showing that SLICER can detect branches and bubbles. (a) Three simulated genes showing the bubble structure. (b) Geodesic entropy computed for the trajectory (top) and recursively for the longest branch (bottom). The dotted line in each plot represents an entropy of 1, which indicates the beginning of a branch. (c) LLE embedding with branches colored. Black is the initial path that splits into two branches (red and blue). The shorter arm of the initial branch then branches again (yellow and green) at the end of the bubble. (d) Plot showing the boundaries of the bubble (blue) as detected by SLICER.

The geodesic entropy profile of this simulated dataset (top graph in Fig. 3b) shows a spike followed by a drop and then another spike. The first position at which the geodesic entropy exceeds 1 indicates the branch in the trajectory. The points colored red in Fig. 3c correspond to one branch, and the points colored blue, yellow, and green correspond to the other. We then recursively computed geodesic entropy on the longer of the two branches. The recursive geodesic entropy profile (bottom of Fig. 3b) indicates that the initial branch gives rise to another branch (yellow and green points in Fig. 3c). The second branch arises because one side of the bubble is slightly shorter than the other, causing the shortest paths from the start point to wrap around past the end of the bubble to reach the end of the red branch. The location of the second branch thus indicates the true end of the bubble. Fig. 3d shows the bubble correctly identified by SLICER colored in blue.

We also tested the robustness of SLICER’s branch detection in the presence of increasing noise and proportion of irrelevant genes (Fig. S4). We used the percentage of cells assigned to the correct branch as a metric for the accuracy of branch detection. This analysis showed that SLICER is able to identify the correct branch assignment for cells even in the presence of irrelevant genes (Fig. S4b) and noise (Fig. S4c), although it appears that noise affects the branch assignment more than irrelevant genes.

### Developing Mouse Lung Cells

We next ran SLICER on previously published data from developing mouse lung cells [6]. The data were generated as follows: cells from the developing bronchio-alveolar epithelium were extracted from embryonic mice on days E14.5, E16.5, and E18.5. The developing lung epithelium during this stage of development contains progenitor cells, intermediates, and cells committed to one of two specialized cell fates (Alveolar Type 1, AT1 and Alveolar Type 2, AT2) [7]. AT2 cells from adult mice (postnatal day 107) were also extracted and sequenced for comparison. We computed gene expression levels using RSEM v. 1.2.8 and the UCSC mm 10 gene annotations. Cells with less than 1000 genes detected at or above 1 FPKM were omitted from further analysis, leaving 183 out of 198 cells. We then log-transformed the expression levels but did not filter the genes in any way.

Each cell in this dataset represents a “snapshot” observation of the sequential process of gene expression changes required for differentiation. Our goal is to investigate the precise sequence of changes, which are not completely understood, although some marker genes for the AT1 and AT2 cell types are known. Differentiation may proceed at different rates across the set of cells, necessitating the use of an internal clock for monitoring differentiation progress, rather than relying strictly on the time point. We therefore would like to construct a trajectory that captures the sequential relationships among the cells undergoing the differentiation process. In addition, this dataset represents an excellent test for the branch detection capabilities of SLICER, because the cells are differentiating toward one of two cell fates, each with a handful of known marker genes.

To determine a set of genes to use in building the trajectory, we selected genes whose expression level variance exceeded their “neighborhood variance” (see Methods). This method produced a list of 660 genes. We next computed a 2D embedding of the data using LLE (Fig. 4a). We then picked a starting cell, constructed a nearest neighbor graph in the low-dimensional space, and found single-source shortest paths from the starting cell using Dijkstra’s algorithm.

**Figure 4.**
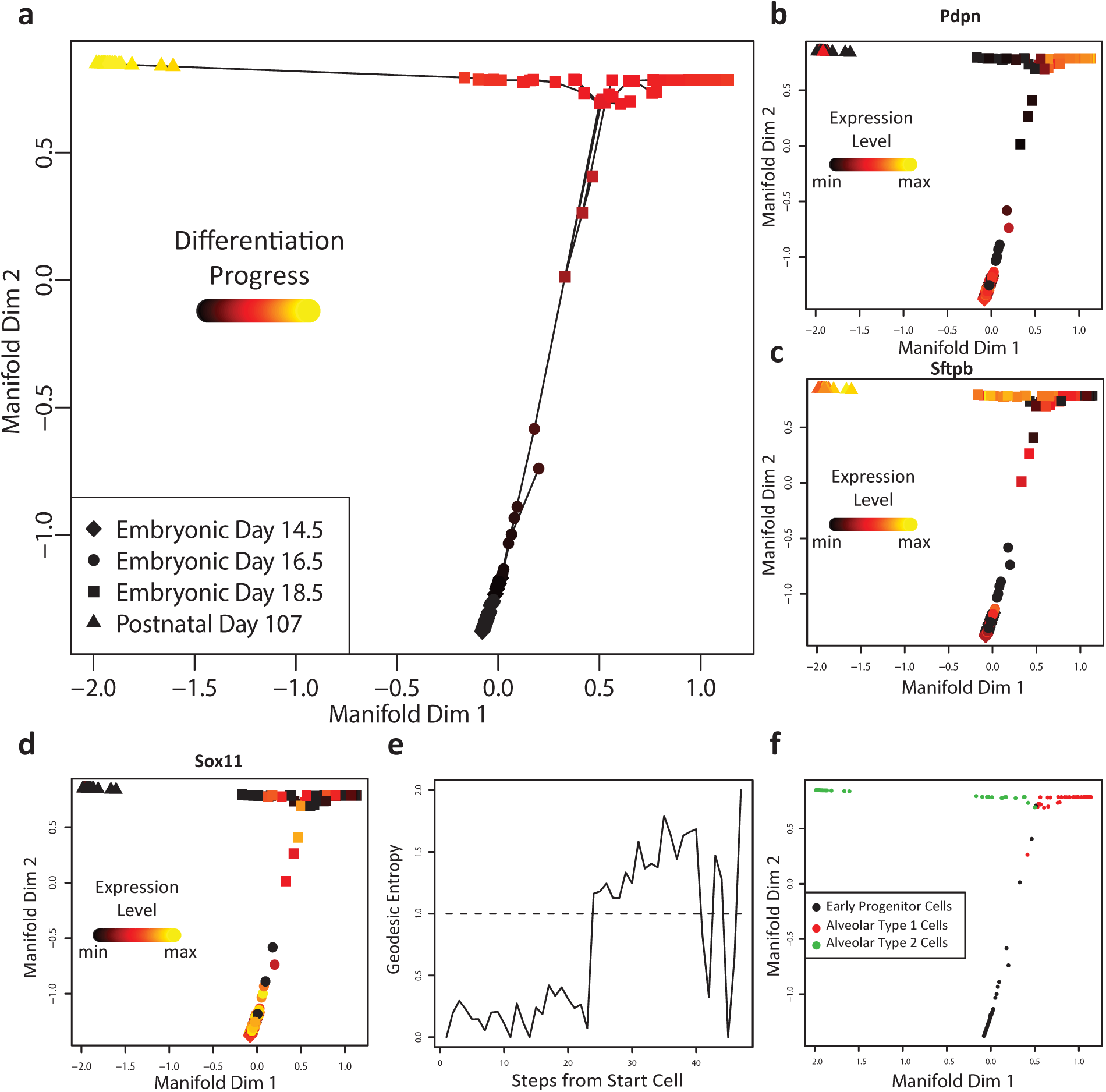
SLICER applied to cells from the developing mouse lung. (a) Cellular trajectory inferred by SLICER. The shape of each point indicates the time point (note that this information is used only after the fact for assessing whether the trajectory makes sense, not for constructing it). Color corresponds to inferred geodesic distance from the start cell (“differentiation progress”). The lines indicate edges used in the shortest paths to each point. Panels (b) through (d) show the expression levels of marker genes in each cell, with the cells ordered by developmental time. (b) shows a marker for alveolar type 1 cells. (c) is an alveolar type 2 marker and (d) is a marker for early progenitor cells. (e) Geodesic entropy plot for the trajectory shown in panel (a). The dotted line represents an entropy value of 1, the threshold for branch detection. (f) Cells colored according to the branches that SLICER assigned using geodesic entropy. Note that no annotations were used in assigning cells to branches; instead, the interpretations indicated in the legend (AT1, AT2, or EP) were deduced based on marker genes such as those shown in panels (b)-(d) after branch assignment.

As Fig. 4a shows, the trajectory reconstructed by SLICER places cells in an order that is clearly related to the day of development. Based on the labels indicating the days on which the cells were extracted, starting at the bottom of the figure and moving to the top and then left or right, seems to correspond to progress through development. In this ordering, the cells separate well by day of development. However, there are some exceptions: cells from days E14.5 and E16.5 overlap significantly, indicating that few changes occur during that two-day period. In contrast, there is a wide separation between day E18.5 and the fully differentiated AT2 cells from post-natal day 107. Another salient feature of the SLICER trajectory is that there appears to be a branch, with some cells positioned to the left of the early progenitors approaching the AT2 cells and some to the right of the early progenitors.

To further investigate the trajectory inferred by SLICER, we examined the expression levels of several genes that were previously validated [6] as markers of mouse lung development (Fig. 4b-d and Supplementary Figure 5). The AT1 marker gene *Pdpn* should show moderate expression in early progenitor cells, high expression in AT1 cells, and low expression in AT2 cells [6]. As Fig. 4b shows, *Pdpn* expression gradually increases along the continuum from early progenitor cells to AT1 cells, matching the expected pattern. Similarly, the AT2 marker *Sftpb* shows increasing expression moving along the trajectory from early progenitors to adult AT2 cells but not AT1 cells (Fig. 4c). Additionally, the transcription factor *Soxll*, which plays a role in tissue remodeling during early lung development [6, 8], shows decreasing expression levels with increasing distance from the start of the trajectory (Fig. 4d). Collectively, the expression patterns of *Pdpn, Sftpb*, and *Soxll* confirm that the SLICER trajectory represents a continuum of cells ordered by differentiation progress from early progenitor cells to either AT1 or AT2 cells.

We also used the branch detection capability of SLICER to infer the presence and location of a branch in the differentiation process. Approximately 25 steps from the starting cell, the geodesic entropy of the trajectory exceeds 1, indicating the beginning of a branch (Fig. 4e). Based on the above investigation of known marker genes, this location appears to represent a decision point for a differentiating cell, after which a cell proceeds toward either the AT1 or AT2 cell fate. After detecting the existence and location of a branch in the trajectory, we used SLICER to assign each cell to a branch (Fig. 4f).

### Mouse Neural Stem Cells

We ran SLICER on previously published data from mouse adult neural stem cells [4]. In this study, cells were harvested from the subventricular zones of adult mice with the goal of determining how gene expression changes during neural stem cell activation after a brain injury [4]. Only one cell fell below the cutoff of 1000 genes detected, leaving 271 out of 272 cells.

We again selected genes by comparing sample variance and neighborhood variance. This yielded a list of 661 genes. Figure 5a shows the resulting trajectory. The embedding has a clear trajectory-like shape, with most of the cells lying along a horizontal path. There are also two clusters of cells, one close to the main group of cells along the horizontal axis, and one in the upper right corner of the plot.

**Figure 5.**
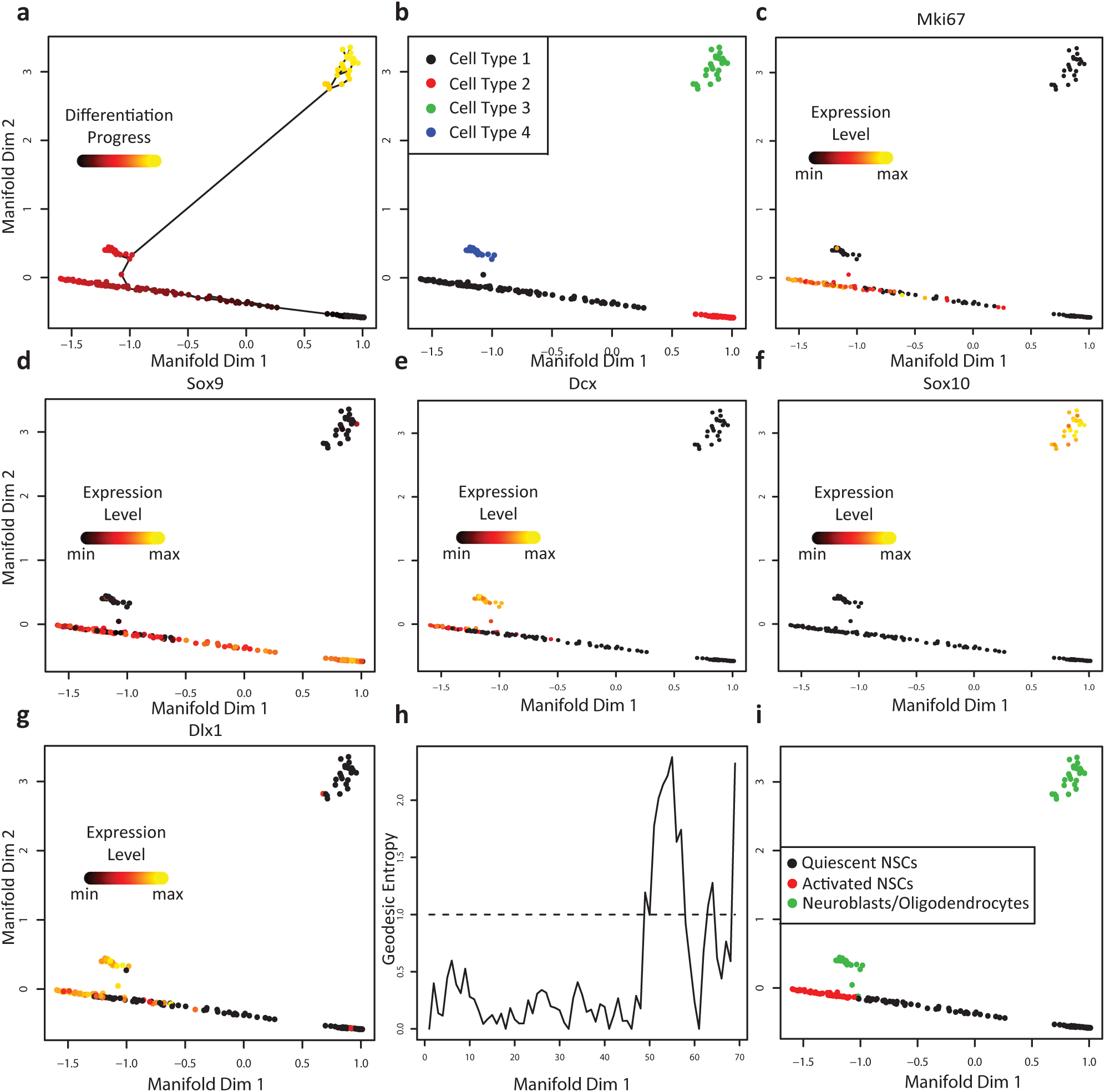
SLICER applied to mouse neural stem cells. (a) Cellular trajectory inferred by SLICER. Color corresponds to inferred geodesic distance from the start cell (“differentiation progress”). The lines indicate edges used in the shortest paths to each point. (b) Clustering using the connected components in the low-dimensional *k*-nearest neighbor graph before trajectory construction identifies four cell types. SLICER provides the option to select which cell types to include when building a trajectory. Panels (c) through (g) show the expression levels of marker genes for different cell types: (c) active neural stem cells. (d) quiescent neural stem cells. (e) neuroblasts. (f) oligodendrocytes, and (g) neuroblasts. (h) Geodesic entropy plot for the trajectory shown in panel (a). The dotted line represents an entropy value of 1, the threshold for branch detection. (i) Cells colored according to the branches that SLICER assigned using geodesic entropy. The interpretations indicated in the legend were deduced based on marker genes such as those shown in panels (c)-(g) after branch assignment.

SLICER has the ability to detect such clusters directly from the low-dimensional *k*-nearest neighbor graph, allowing the user to include or omit certain cell types from trajectory construction (Fig. 5b). For example, in the initial analysis of this dataset, the authors discovered the presence of oligodendrocytes, mature neural cells that were extracted at low levels due to overlap with the markers used to isolate neural stem cells. Based on our analysis of marker genes distinguishing oligodendrocytes and neural stem cells (see below), the green cell type in Fig. 5b corresponds to oligodendrocytes. SLICER thus gives the ability to easily exclude oligodendrocytes from further analysis, although we chose to retain them because they provide a good example of a trajectory with multiple branches (see below).

To investigate whether the trajectory produced by SLICER is related to the activation of neural stem cells, we examined the expression of known marker genes (Fig. 5c-g and Supplementary Figure 6). The *Mki67* gene was previously shown to be a marker for active neural stem cells (aNSCs), and the transcription factor Sox9 is associated with quiescent neural stem cells (qNSCs) [4]. When we colored the trajectory with the expression levels of these marker genes, we found that cells along the x-axis in Fig. 5a show gradual variation, with high qNSC marker expression on the right and high aNSC marker expression on the left (Fig. 5c-d). This suggests that these cells represent a continuum of states from quiescent to active neural stem cells. The expression of *Dcx*, a neuroblast marker that is also responsible for the proper migration of differentiating neurons [4, 9], is expressed at high levels in the cluster of cells near the horizontal axis, indicating that this cluster of cells corresponds to neuroblasts (Fig. 5e). The cluster of cells that is far removed from the others shows high expression of the oligodendrocyte transcription factor *Sox10* [4], indicating that these cells are oligodendrocytes (Fig. 5f). The *Dlx1* gene encodes a transcription neuroblast-associated transcription factor, and it was observed in [4] that some of the aNSCs also expressed this marker, indicating the initiation of a differentiation program in the aNSCs. Our analysis confirms this result (Fig. 5g).

One of the key advantages of SLICER is the ability to identify multiple levels of branches automatically using geodesic entropy, as the synthetic data example in Fig. 3 showed. The neural stem cell dataset provides an excellent opportunity to demonstrate this capability on real data because of the presence of three distinct cell types. The geodesic entropy profile of the trajectory indicates a branch about 50 steps from the starting cell. This branching event corresponds to the distinction between aNSCs and neuroblasts (Fig. 5h-i). We next computed geodesic entropy recursively on each of the top-level branches identified (red and green cells shown in Fig. 5i). SLICER identified a second branch separating neuroblasts from oligodendrocytes but did not detect a branch in the aNSCs (Supplementary Figure 7).

Because the cost of single cell RNA-seq depends strongly on the number of cells to be sequenced, the number of cells required to construct a trajectory is an important question. However, the number of cells needed depends strongly on the biological process under consideration. Factors such as the number of branches, relative size of each branch, and extent of the changes across the sampled set of cells all can affect this number. With these caveats in mind, we have addressed this question by investigating, for both of our biological datasets, how much the trajectory changes when SLICER is given a random subset of the cells rather than the full dataset (Fig. S8). The results indicate that, for both datasets, the ordering of the cells is relatively stable even with as few as 20% of the cells. The assignment of cells to branches is stable down to 20% of the cells for the distal lung epithelium dataset, but the assignment accuracy steadily declines for the neural stem cell dataset. The reason for this difference is most likely that there are more branches in the neural stem cell dataset, and a smaller proportion of cells occur after the branch points. In contrast, the single branching event in the lung dataset is roughly an even split and occurs mid-way through the trajectory. Thus, in this case the separation between the cell fates is maintained even when only a few cells are used to build the trajectory.

### Comparison with Other Methods

In order to assess the performance of SLICER in relation to other approaches, we ran ICA and Wanderlust on the lung and neural stem cell data and compared the results from all three approaches. We used the set of genes selected by SLICER to ensure that the results from all three approaches were directly comparable. We also set the number of nearest neighbors for Wanderlust to the same values used by SLICER.

The ICA embedding of the mouse lung data in Fig. 6a resembles the trajectory inferred by SLICER, detecting a single main path with a prominent branch. However, the arrangement of the points in the embedding is noticeably more diffuse and less “trajectory-like” than the SLICER result shown in Fig. 4. In addition, the geometric relationship between early progenitor cells and AT2 cells is somewhat different than that inferred by SLICER (compare Fig. 4a and Fig. 6a). It appears that tracing a shortest path from early progenitor cells to AT1 cells in Fig. 6a would pass through AT2 cells, while the SLICER branching analysis and marker gene expression suggest that these cells should fall on different branches. The ICA embedding of the neural stem cells shows a similar overall shape to the SLICER trajectory (compare Fig. 5a and Fig. 6b). Once again, however, the overall shape of the ICA embedding is much more amorphous, and an ordering of the cells from quiescent to active is much less apparent than in the SLICER trajectory shown in Fig. 5a.

**Figure 6.**
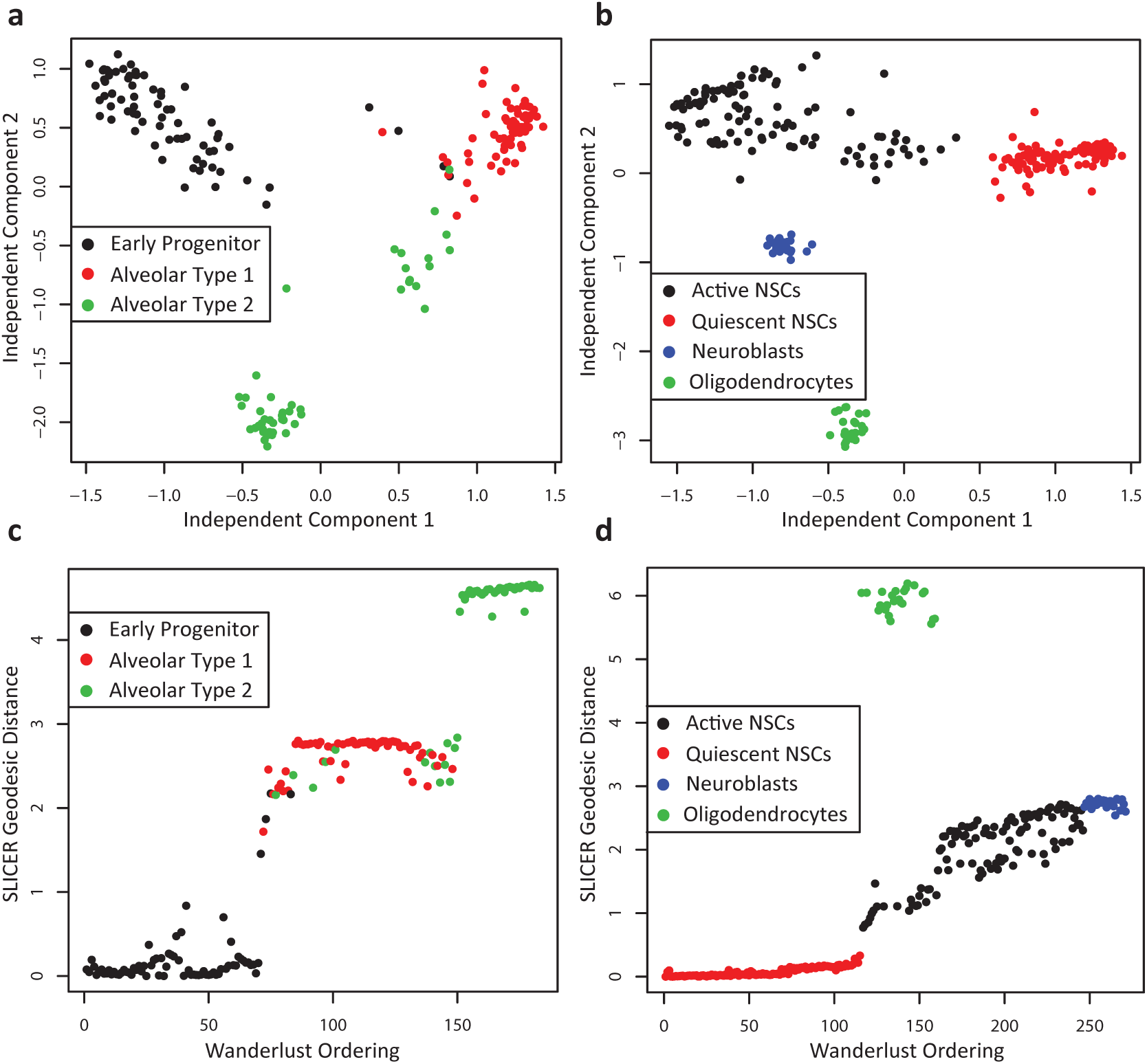
ICA and Wanderlust results from mouse lung and neural cells. Note that the genes selected by SLICER were used as input to both ICA and Wanderlust to ensure an accurate side-by-side comparison. (a) ICA embedding of mouse lung cells. The colors correspond to the branch assignments from SLICER. (b) ICA embedding of mouse lung cells. Colors correspond to the SLICER cell type assignments from Fig. 5b. (c) Comparison of one-dimensional Wanderlust ordering (x-axis) and SLICER geodesic distance (y-axis) for mouse lung cells, (d) Comparison of one-dimensional Wanderlust ordering (x-axis) and SLICER geodesic distance (y-axis) for mouse neural cells.

Because Wanderlust produces only a one-dimensional ordering of cells rather than a two-dimensional embedding, we plotted the Wanderlust ordering of cells against the SLICER geodesic distance (Fig. 6c-d). The two tools agree on the relative ordering of mouse lung cell types, with early progenitor cells preceding AT1 cells and most AT2 cells (Fig. 6c). However, it is important to note that because Wanderlust assumes that a trajectory does not branch, the Wanderlust ordering suggests that lung differentiation process moves from early progenitor cells to AT1 cells, then AT2 cells. In addition to obscuring the true sequence of events in the differentiation process, the existence of multiple cell fates is lost in this approach, underscoring the importance of detecting branches in a trajectory. The Wanderlust ordering of neural stem cells agrees with SLICER on the relative ordering of qNSCs, aNSCs, and neuroblasts (Fig. 6d). One exception to note, however, is that Wanderlust places the oligodendrocytes in the middle of the ordering, interleaving them with aNSCs.

## Discussion

We developed SLICER (Selective Locally linear Inference of Cellular Expression Relationships), a method for inferring cellular trajectories from single cell RNA-seq data. The key advantages of our approach are (1) the ability to discover which genes to use in building the trajectory without prior knowledge of the process, (2) a novel method for detecting the number and location of branches and bubbles, and (3) the ability to infer nonlinear trajectories. In addition, our evaluation of SLICER on synthetic data shows that the method is highly robust to the presence of genes unrelated to the process. Our simulations also show the importance of modeling nonlinear behavior. It is worth noting that the choices of highly nonlinear functions such as cosine and square root that we used to simulate gene expression levels are not an unrealistic representation of biological systems, given such phenomena as cell-cycle genes and Michaelis-Menten enzyme kinetics [10].

We showed that SLICER can detect branches and bubbles in simulated trajectories. In addition to making the trajectory construction process simpler for the user by eliminating the need for manually counting the number of branches, our branch detection approach provides two important advantages. First, it can be used to detect branches even if the number of manifold dimensions is greater than two or three. For example, a recent paper that examined a hematopoiesis differentiation continuum used a four-dimensional projection to construct a cell trajectory [11]. The method of manual inspection cannot be used directly in this case, because there is no easy way to visualize a four-dimensional space. Second, geodesic entropy is a metric that characterizes the degeneracy of a trajectory, which opens the door to developing a statistical model of branch significance. Such a model, although beyond the scope of the current paper, would be very useful for detecting branching events corresponding to rare cell types or rare alternate outcomes of a process.

Our branch detection approach is capable of identifying loops or “bubbles”, but to the best of our knowledge, no “bubbles” have yet been discovered through analysis of real single cell RNA-seq data. However, it is interesting to consider what might cause a bubble, and what might be the biological significance of such a geometry. One situation in which a bubble might arise is when there are multiple possible sequences of events that can lead a cell to the same position in a process. For instance, there may be cases in which the master regulator genes A, B, and C must all be turned on in order to complete some process, but it does not matter which of the genes is turned on first. If each of these three genes subsequently induces a separate regulatory cascade, cells could reach a final end state through several distinct sequences of gene expression changes. Another way in which a bubble might occur is if cells within a single initial population receive distinct signals that eventually lead the cells to a common state. It is possible that detecting a bubble from real data would require a process to be well-sampled, which means that many cells would be required. As increasingly high-throughput techniques for single cell isolation and sequencing emerge [12, 13], the number of cells sequenced is likely to increase dramatically. We hypothesize that as more and more biological processes are studied at the single cell level with increasing numbers of cells, examples of different processes arriving at the same outcome will be discovered.

## Methods

### Trajectory Reconstruction

We use locally linear embedding (LLE) [14], a nonlinear dimensionality reduction technique, to reconstruct cellular trajectories. LLE belongs to the class of nonlinear dimensionality reduction techniques, which includes a number of methods, such as Isomap [15]. Hessian LLE [16]. Laplacian eigenmaps [17], and diffusion maps [18]. Nonlinear dimensionality reduction techniques have been widely on high-dimensional data used to perform denoising and feature extraction for subsequent classification or regression. For example, such techniques were used to estimate head pose angle and age from images of human faces [19, 20]. We initially experimented with Isomap, Hessian LLE, Laplacian eigenmaps, and diffusion maps and found that LLE seemed to give the best results.

To infer a trajectory using LLE, we take as input a matrix of expression levels with *n* samples and m genes *E*_*n*×*m*_ = (*e*_*ij*_), where *e*_*ij*_ is the expression of gene *j* in sample *i*. Then we perform LLE on *E*_*n*×*m*_ to give a low-dimensional embedding *L*_*n*×*d*_. Our analysis of synthetic and real datasets indicates that *d* = 2 is a reasonable choice (see below for more detailed discussion of this point). LLE performs dimensionality reduction in two steps. First, a set of reconstruction weights *W*_*n*×*k*_ is learned so that each point in high-dimensional space is represented as a linear combination of its *k*-nearest neighbors, where *k* is a chosen constant:

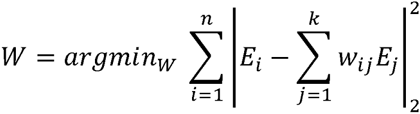

The row sums of W are constrained to 1 to ensure translational invariance [14]. Then, the weights are used to solve for the coordinates of each point in *d*-dimensional space:

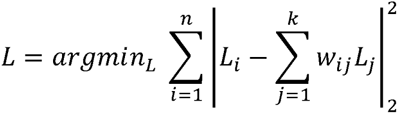

The sum-to-one constraint on the reconstruction weights and the form of the weight equations ensure that the low-dimensional reconstruction preserves the high-dimensional geometry of the points [14].

After embedding the data using LLE, we build a *k*-nearest neighbor graph in the low-dimensional space. Then we use Dijkstra’s algorithm [21] to find the single source shortest paths from a user-specified start point. These shortest paths can be thought of as geodesics that characterize the shape of the cell trajectory manifold, and the length of the shortest path to a particular point represents its geodesic distance from the source point. These geodesic distances can then be used to order the points according to their progress through a process.

The question of the best choice for dimensionality (*d*) is difficult to answer for the trajectory construction problem, because the ground truth cell ordering is unknown for the biological data, and the synthetic data are generated to yield a specific intrinsic dimensionality. While developing SLICER, we explored using intrinsic dimensionality estimators such as packing numbers [22] and nearest neighbor estimation [23] to determine *d*, but tests on our synthetic data showed these methods to be unreliable and highly sensitive to noise. In addition, most of these methods require setting a scale parameter, which simply moves the problem of choosing the dimensionality parameter back one level. Most cell trajectory studies to date have used *d* = 2, and this seems to yield biologically meaningful results. To our knowledge, only one study has used *d* > 2 [11]. For the datasets that we used here, *d* = 1 will hide any branches in the trajectory (see Fig. 6), and *d* = 3 produces an embedding that is not qualitatively different than *d* = 2 (Fig. S9). We note that SLICER allows the user to specify the number of dimensions, and works for *d* ≥ 2.

### Gene Selection

Selecting the genes to use when constructing a trajectory is a key step in the process. Both Monocle and Wanderlust require the pre-selection of genes based on some sort of prior knowledge. The Monocle paper selected genes that exhibited differential expression in bulk RNA-seq samples taken from the initial and final time points. However, in some cases, cells are collected at only a single time point, and furthermore it would be ideal to have a method for selecting genes without the need for prior knowledge provided by additional experiments.

We developed an approach for selecting genes based on a simple intuition: If a gene is involved in progression along a cellular trajectory, we expect to see gradual changes in the expression of the gene along the trajectory. Conversely, if a gene is not involved in the sequential progression, the gene should fluctuate in a manner independent of the trajectory. Because gene selection must be performed before trajectory construction, selecting genes directly based on whether they are related to the trajectory is not possible. Instead, we note that points close together in Euclidean space are likely to lie close together on the manifold, and we can thus use the similarity of genes in neighboring points to approximate the change in a gene moving along the trajectory. Specifically, for a gene *g*, we calculate the sample variance 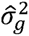 of *g* across all samples. Then, we compute the “neighborhood variance”

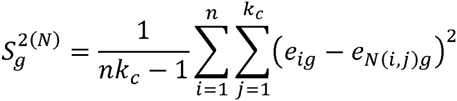

where *e*_*ij*_-is the expression level of the *j*th gene in the *i*th sample, *N*(*i*,*j*) is the *j*th nearest neighbor of sample *i*, and *k*_*c*_ is the minimum number of neighbors needed to yield a connected graph. Intuitively, the quantity 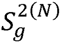 is like a sample variance computed with respect to neighboring points rather than the mean, and it measures how much *g* varies across neighboring samples. To select the genes that are most likely to be involved in the trajectory, we pick *g* such that 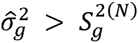. These are genes that show more gradual variation across neighboring points than at global scale.

In biological datasets, genes often cluster into co-expressed modules, so an important question is how our gene selection method handles co-expressed genes. Because the variance and “neighborhood variance” are computed for each gene separately, genes related to the trajectory will be selected whether or not they are co-expressed. Conversely, genes that are unrelated to the trajectory will not be selected even if they are co-expressed. Examining the correlation matrix of selected genes from the two biological datasets shows that there is a high degree of co-expression, with genes clustering into co-expressed modules (Fig. S10). We also note that our simulations include “genes” that show strong co-expression because they are generated from a handful of functions simulating shared gene regulatory mechanisms. Our simulation results indicate that the gene selection approach works well for these co-expressed genes (Fig. 2, Fig. S3, Fig. S4).

### Choosing the Number of Neighbors

Previous approaches for selecting the number of neighbors for LLE have relied upon similarity metrics comparing the relative distances of points in the full space and the embedded space [24, 25]. We initially tried such approaches and found that they work fairly well on the simulated data but tend to recommend improbably large values for *k* when run on real data. Consequently, we developed an alternate method that is tailored to the particular manifold shape that we expect to see in this problem. In particular, we expect a trajectory to resemble a long, narrow shape rather than an amorphous point cloud.

To formalize this intuition, we use the notion of alpha convex hull [26]. The *α*-hull of a set of points is the intersection of all closed discs with radius *α* that contain all of the points. For a given *k*, we perform LLE and compute the length *I* of the longest shortest path (see Trajectory Reconstruction). We then find the area *α* of the *α*-hull with *α* = 1/10. This choice of a corresponds to the fraction of the length that contains roughly 10% of the datapoints. Using the area of the *α*-hull allows us to compute the “width” of the embedding: *w*_*k*_ = *a*/*l*. The quantity *w*_*k*_ quantifies how much the embedding resembles a trajectory, and we choose *k* = *argmin*_*k*_{*w*_*k*_}. Supplementary Figure 1 shows an example of the longest shortest path and *α*-hull for a 2D LLE embedding.

### Detecting Branches

In some cases, the manifold describing a cellular trajectory possesses important properties such as branches. For example, the Monocle paper found a branch in the trajectory corresponding to a split in development resulting in two different cell fates [1]. We developed a novel approach for characterizing the branching structure of a manifold. Our approach can detect the location and number of branches. In addition, we can readily distinguish branches from convergences and bubbles. To do this, we take as input the set of shortest paths used to characterize the trajectory (see Trajectory Reconstruction) and use them to compute a metric that we term geodesic entropy. Intuitively, our approach lines up the shortest paths from the start point to all other points and asks whether the paths use similar vertices. Let *t*_*i*_ = {*s* = *v*_1_,… *v*_*k*_,… *v*_*l*_ = *i*} be the shortest path along the manifold from the starting point *s* to point *i* that passes through the *l* points *v*_1_,…,*v*_*k*_,…,*v*_*l*_. Denote the *k*th vertex on the shortest path from *s* to *i* by *t*_*i*_(*k*). Consider the set *S* of shortest paths to each point on the manifold, then

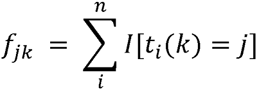

is the number of these paths that pass through point *j* k at distance *k*, where *I*[·] is an indicator function. The fraction of all paths in *S* that pass through vertex *j* at distance *k* is

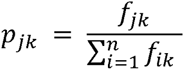

Finally we define *H*_*k*_ as the Shannon entropy of *p*_*k*_:

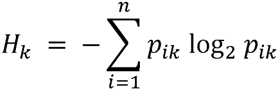

We refer to the quantity *H*_*k*_ as *geodesic entropy* because it describes the vertex composition degeneracy of the shortest paths along the manifold (geodesics). If most of the paths are similar in the first *k* vertices, then the geodesic entropy *H*_*k*_ will be low (approximately zero), indicating that the manifold does not branch. High geodesic entropy, on the other hand, indicates that multiple distinct vertices are being used along the shortest paths. In fact, following the information theoretic interpretation of entropy as the number of bits needed to transmit a message across a channel, a geodesic entropy of *H*_*k*_ means that there are approximately 2^*H*_*k*_^ distinct paths *k* vertices from the start point.

Supplementary Figure 2 shows an example of a branching trajectory and illustrates how geodesic entropy at *k* = 10 steps from the starting cell is computed for this example. To compute *f*_*ik*_ and *f*_*jk*_, count the number of shortest paths that contain *i* and *j* at position *k*; these numbers are 8 and 9 respectively. This means that the probability of seeing vertex *i* at position *k* is 8/17, and the probability of seeing vertex *j* at position *k* is 9/17. If we treat (*p*_*ik*_, *p*_*jk*_) as a probability distribution, we can take calculate the geodesic entropy to obtain *H*_*k*_ ≈ 1.

We use geodesic entropy to detect the location and number of branches and to assign points to branches as follows. Choose *d* as the smallest value of *k* such that *H*_*k*_ ≥ 1. This represents the number of steps from the start point along the manifold geodesics at which at least two branches are first detected. The approximate number of branches at *d* is given by 2^*H*_*d*_^. Now decrement *d* until you reach a value *c* such that only one value of *p*_*ic*_ is positive (or greater than some *ϵ*; we used *ϵ* = 0.05). This represents a vertex at which there is still only one path but beyond which the branch occurs. Now take *b* = *c* + 1 as the location of the branch and pick the 2^*H*_*d*_^ “distinguishing points” with the highest *p*_*ib*_ values. A point *i* can then be assigned to a branch based on the value of *t*_*ib*_, that is, which of the “distinguishing points” is used at position *b* in the shortest path to *i*. Points with shortest paths containing fewer than *b* vertices fall before the branch. As a practical detail, geodesic entropy will sometimes be high if very few cells are under consideration. For example, at the end of a trajectory, if the shortest path from the start passes through a single cell *c* and ends at each of the *k* neighbors of *c*, the geodesic entropy will be log_*2*_*k* even though there is not really a branch at *c*. This problem can easily be addressed by ignoring any branches with less than some number *t* of cells (SLICER uses *t* = 10 by default).

In addition to detecting branches, geodesic entropy can be used to infer other interesting geometries, such as “bubbles” (see Fig. 3). A bubble is a branch that subsequently converges to a single path and can be detected as a spike in *H*_*k*_ such that points on the distinct branches after the spike are connected downstream of the branch. Complex structures with multiple branches can be unraveled by recursively computing geodesic entropy using the subgraph corresponding to each branch.

We can detect a bubble as follows. We first detect a branch as described above. If the branches identified in this way are connected through the *k*-nearest neighbor graph downstream of the branch point, then this indicates that the branches converge to form a bubble. However, the branches may not be of exactly equal lengths; if they are not, then the shortest paths from the start point will continue past the end of the shorter branch and wrap around the bubble. In such a case, there will be another branch at the end of the bubble, where one set of shortest paths continues around the bubble and the other set exits the bubble (see Fig. 3 for an example of such a case). We can detect this second branch by recursively computing geodesic entropy on the shorter of the two initial branches. The location of the second branch then indicates the end of the bubble. In the case of initial branches that are exactly the same length, the point at which they connect after the initial branch point indicates the end of the bubble.

### Simulating Trajectories

We constructed a set of simulated trajectories to assess the performance of SLICER on inputs with known solutions. To do this, we generated simulated expression levels for genes in such a way that the expression levels are a function of a “process time” parameter *t*. We simulated 5 different “pathways” using distinct families of functions; the genes generated by a single family of functions are analogous to co-regulated genes in a biological pathway that all change in response to a common regulatory mechanism. For the simulations shown in Fig. 2, we used the following 5 functions:

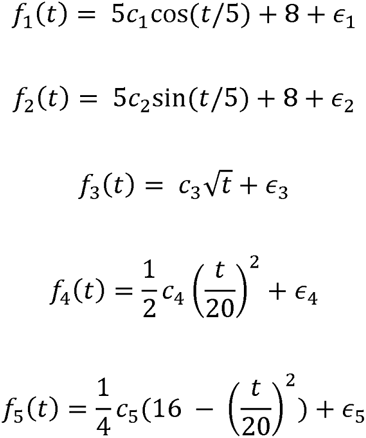

where *c*_*i*_ ~ *N*(1,0.01) and *∈*_*i*_ ~ *N*(0, *σ*^2^). For the simulations in Fig. S3b-d, we used *f*_6_(*t*) = 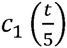, *f*_7_(*t*) = 5 log(*t* + 1) + 8, and *f*_3_, *f*_4_, *f*_5_ as defined above.

The genes used in the simulated are generated by multiplying the value of the corresponding function *f*(*t*) by a normally distributed random variable *c*_*i*_. For the actual values of t, we used the sequence of 801 values 0,0.1,0.2, …,79.9,80. Because each simulated gene depends on *t*, points simulated in this way lie along an essentially one-dimensional manifold (a trajectory) in high-dimensional space. Because in the real data setting we do not know in advance which genes are involved in a trajectory, we also devised a means to simulate genes that are unrelated to the process. To do this, we randomly permute the simulated values of some genes, thus removing their relationship with *t*. The number of such randomly reshuffled genes is controlled by a parameter *p*. As genes are simulated, we pick a set of 5 genes (one from each pathway) to reshuffle. A group of 5 is reshuffled in this way with uniform probability *p*. Randomly permuting the genes (rather than simply sampling from a Gaussian, for instance) ensures that the values lie in the exact same range as the related genes, yet have no relationship with *t*.

To measure the performance of a trajectory reconstruction algorithm, we use the algorithm to produce an ordering of the points, then compare the ordering to the true value of *t* used to generate it. We measure the “percent sortedness” of a list by computing the following quantity:

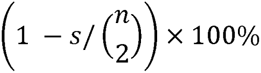

where *s* is the number of pairs of items in the list that are out of order. We chose to use percent sortedness rather than a metric related to distance along the trajectory because dimensionality reduction re-scales the data, which makes it difficult to compare methods that perform dimensionality reduction with those that do not. We used the percentage of points assigned to the correct branch as a metric for evaluating SLICER’s branch detection algorithm.

---

**Supplementary Figure 1:**
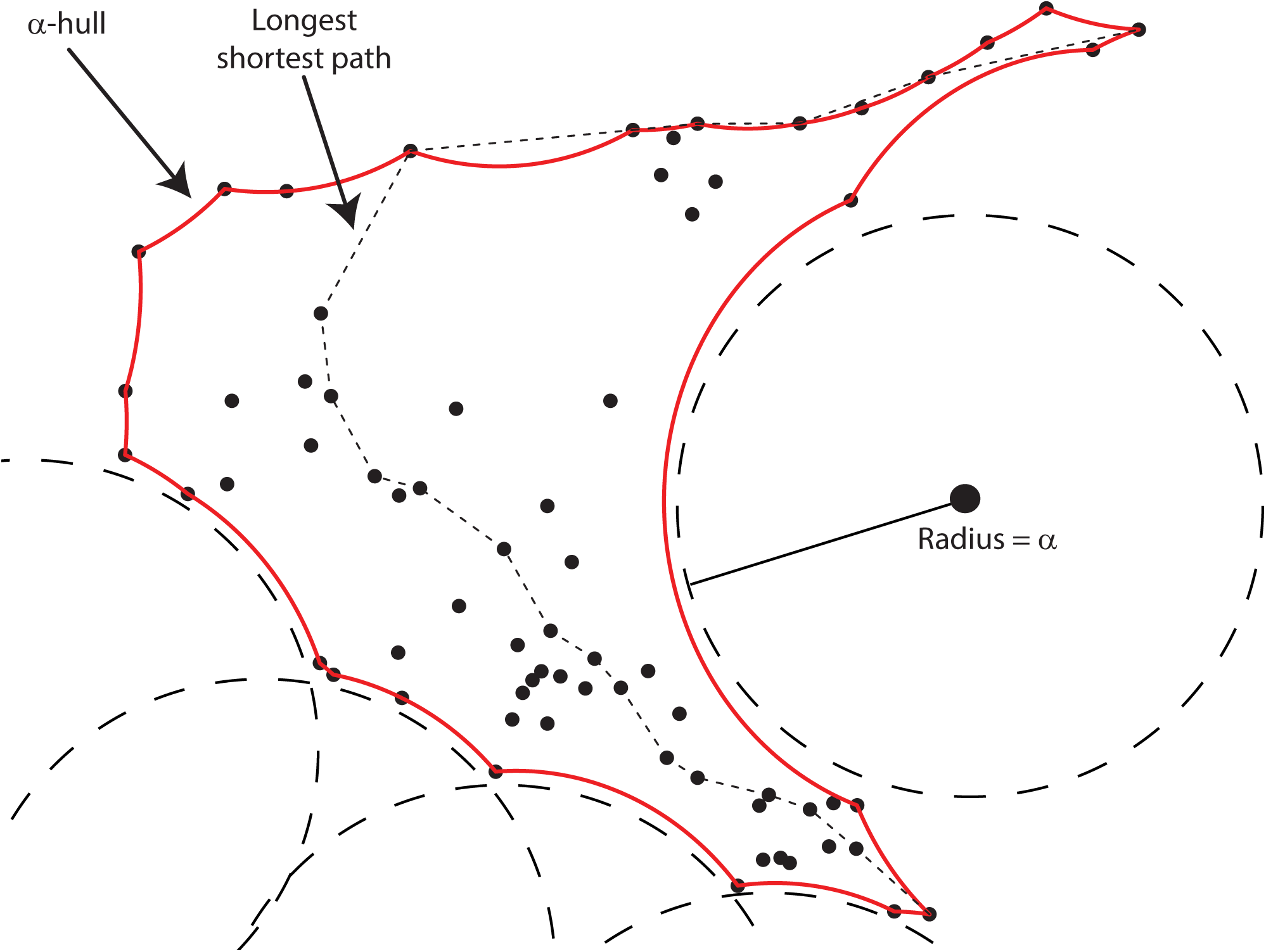
Selecting *k* using the *α*-convex hull. The red border is the alpha-convex hull of the set of points shown, obtained by taking the intersection of the spheres of radius *α* indicated here. The longest shortest path is shown as a dotted line.

**Supplementary Figure 2:**
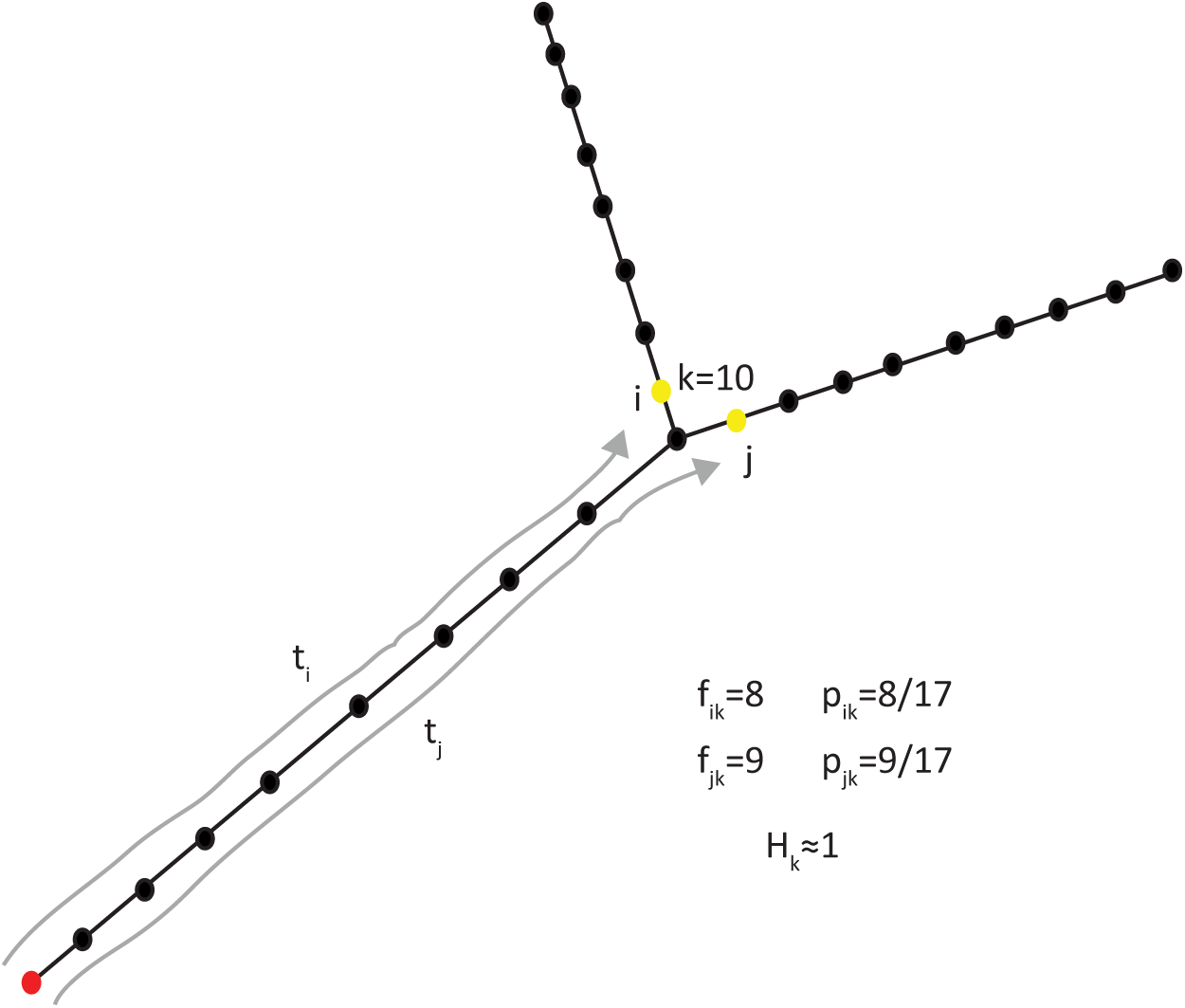
Computing geodesic entropy of a trajectory. The starting cell is indicated in red, two geodesics (shortest paths) are shown in gray, and the cells at *k* = 10 steps away from the starting cell are indicated in yellow.

**Supplementary Figure 3:**
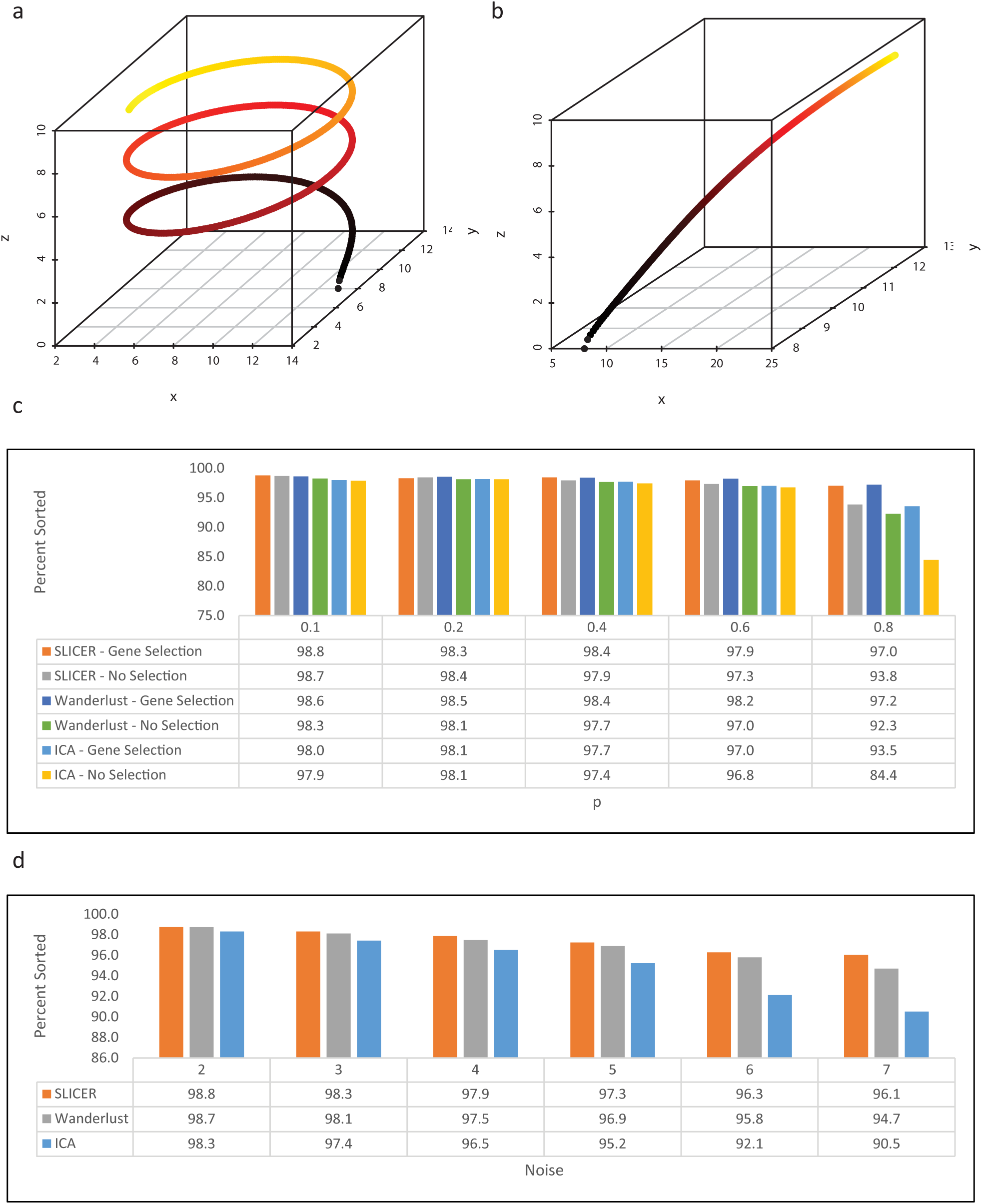
Additional simulations comparing SLICER with other approaches. (a) The first three functions used to generate the synthetic data discussed in Fig. 2. Note the highly curved shape of the trajectory. (b) The first three functions used to generate an additional dataset. This trajectory is much less curved than the one shown in panel a, and ICA thus performs much better on this example. (c) Performance of SLICER and other approaches, with and without gene selection, on the trajectory shown in panel b as the proportion of irrelevant genes increases. Note that the other approaches do not perform gene selection on their own, so the genes selected by SLICER were given as input for this comparison. A noise level of 2 was used for these simulations. Note that the y-axis does not start at 0. (d) Performance of SLICER and other approaches on the trajectory shown in panel b as the noise level increases. To isolate the effect of increasing noise, irrelevant gene proportion of *p* = 0 was used for these datasets. Note that the y-axis does not start at 0.

**Supplementary Figure 4:**
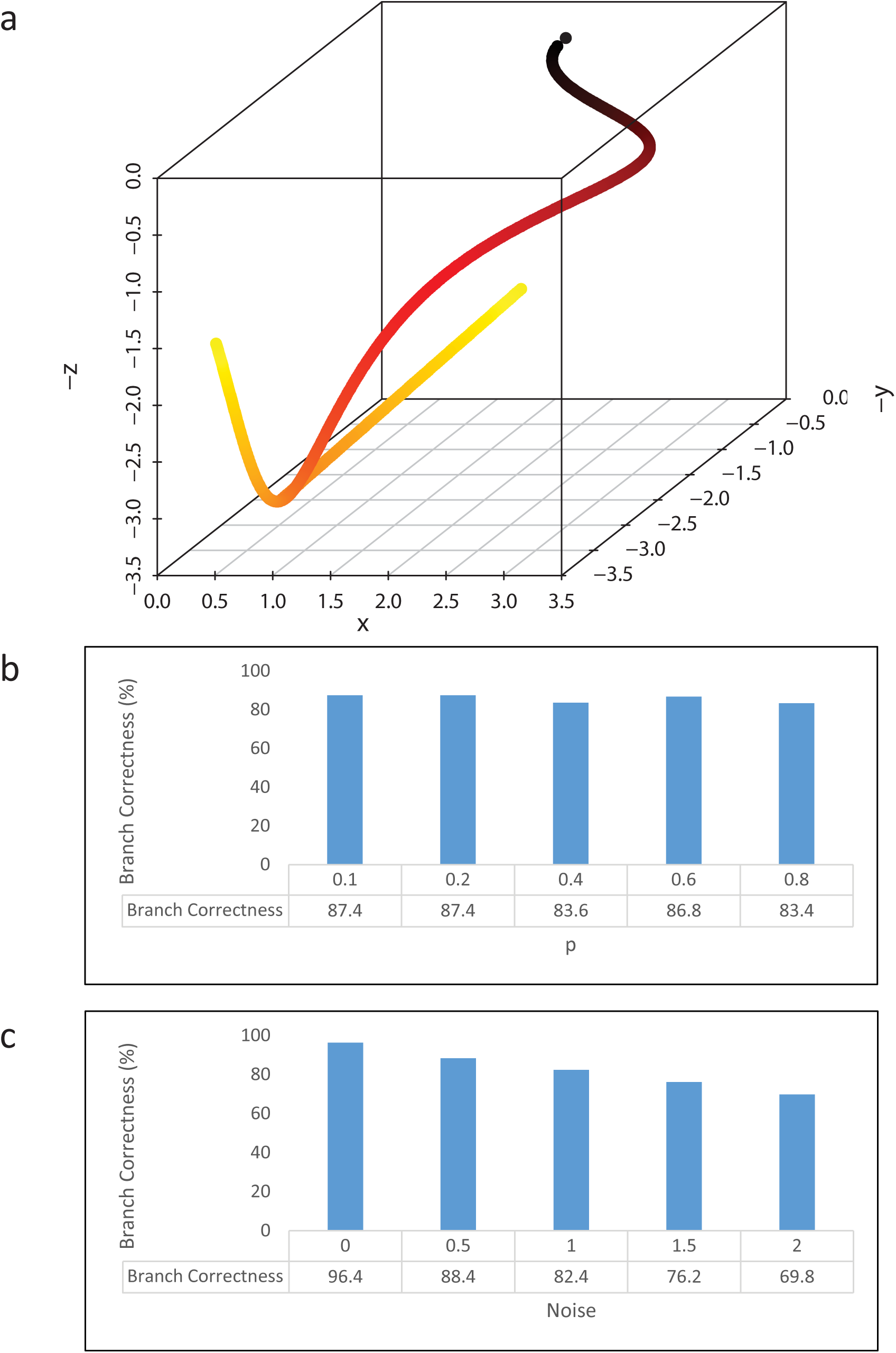
Robustness of branch detection in the presence of noise and irrelevant genes. (a) Simulated branching trajectory used to assess the robustness of SLICER’s branch detection method. (b) Chart showing the percent of cells assigned to the correct branch by SLICER as the proportion of irrelevant genes increases (noise = 0.5). (c) Percent of cells assigned to the correct branch in the presence of increasing noise (*p* = 0).

**Supplementary Figure 5:**
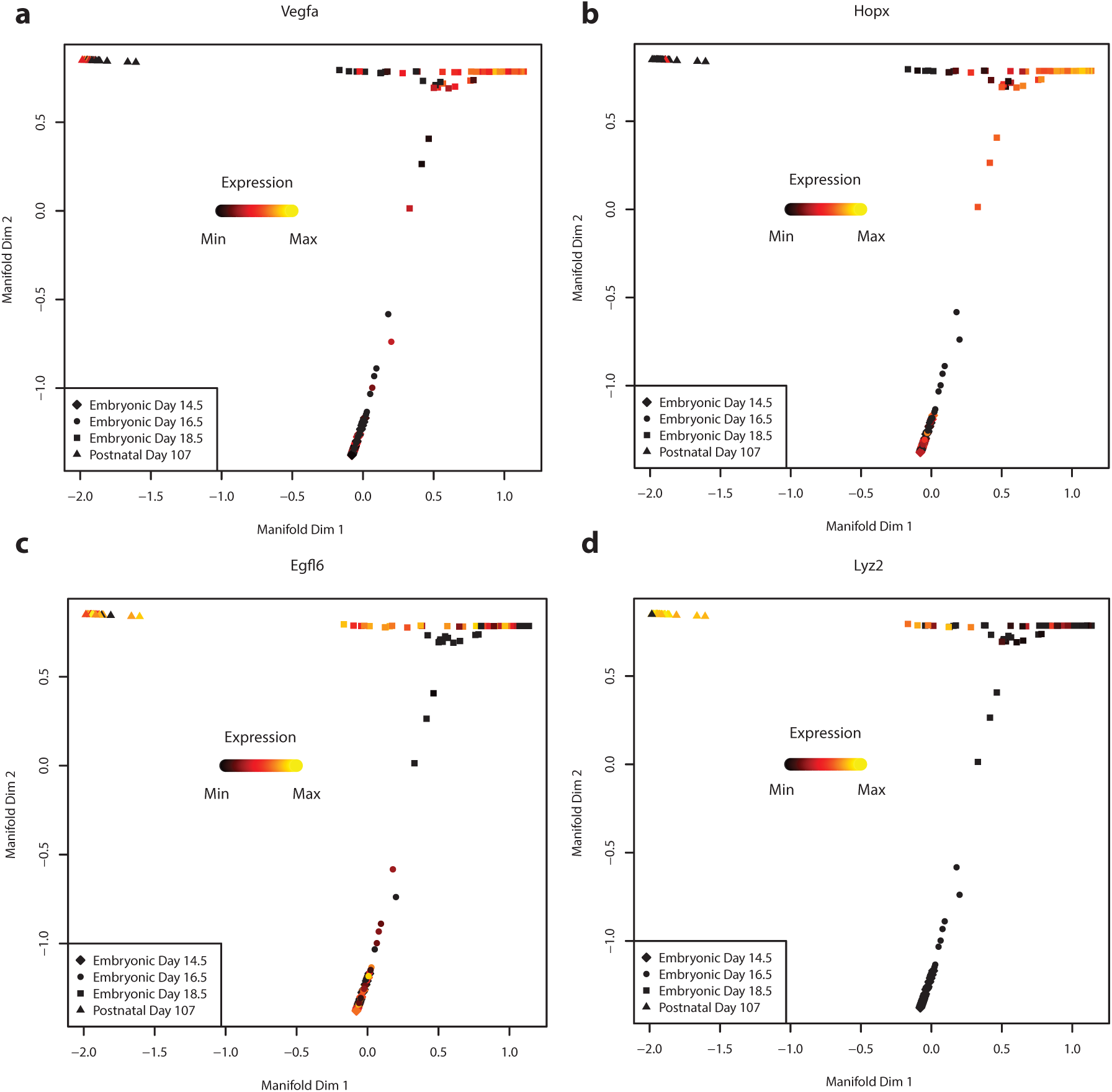
Additional marker genes for mouse lung dataset. (a) and (b) are markers for alveolar type 1 (AT1) cells, (c) and (d) are markers for AT2 cells.

**Supplementary Figure 6:**
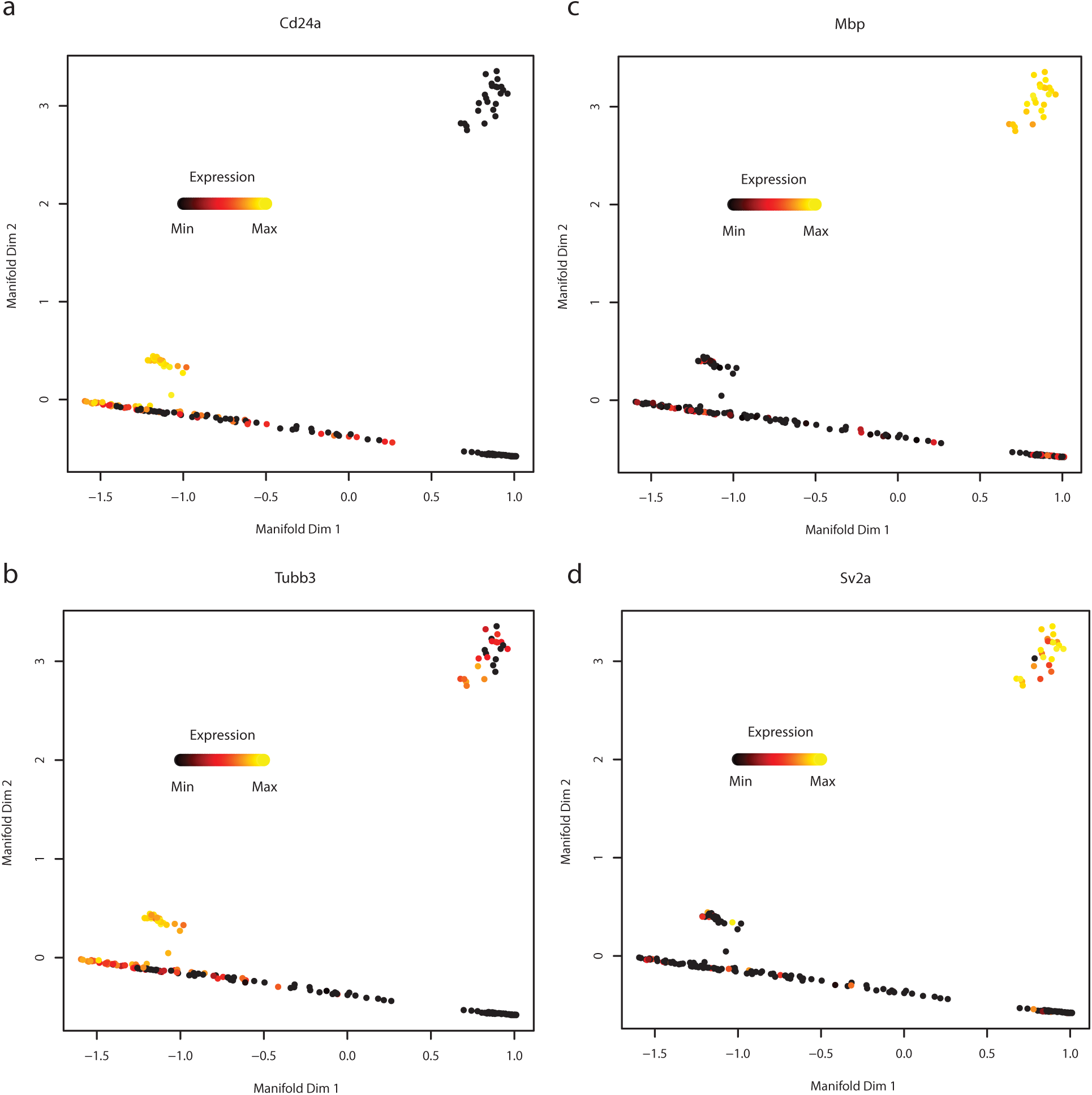
Additional marker genes for mouse neural stem cell dataset. (a) and (b) are neuroblast markers that also show expression in some active NSCs. (c) and (d) are oligodendrocyte markers.

**Supplementary Figure 7:**
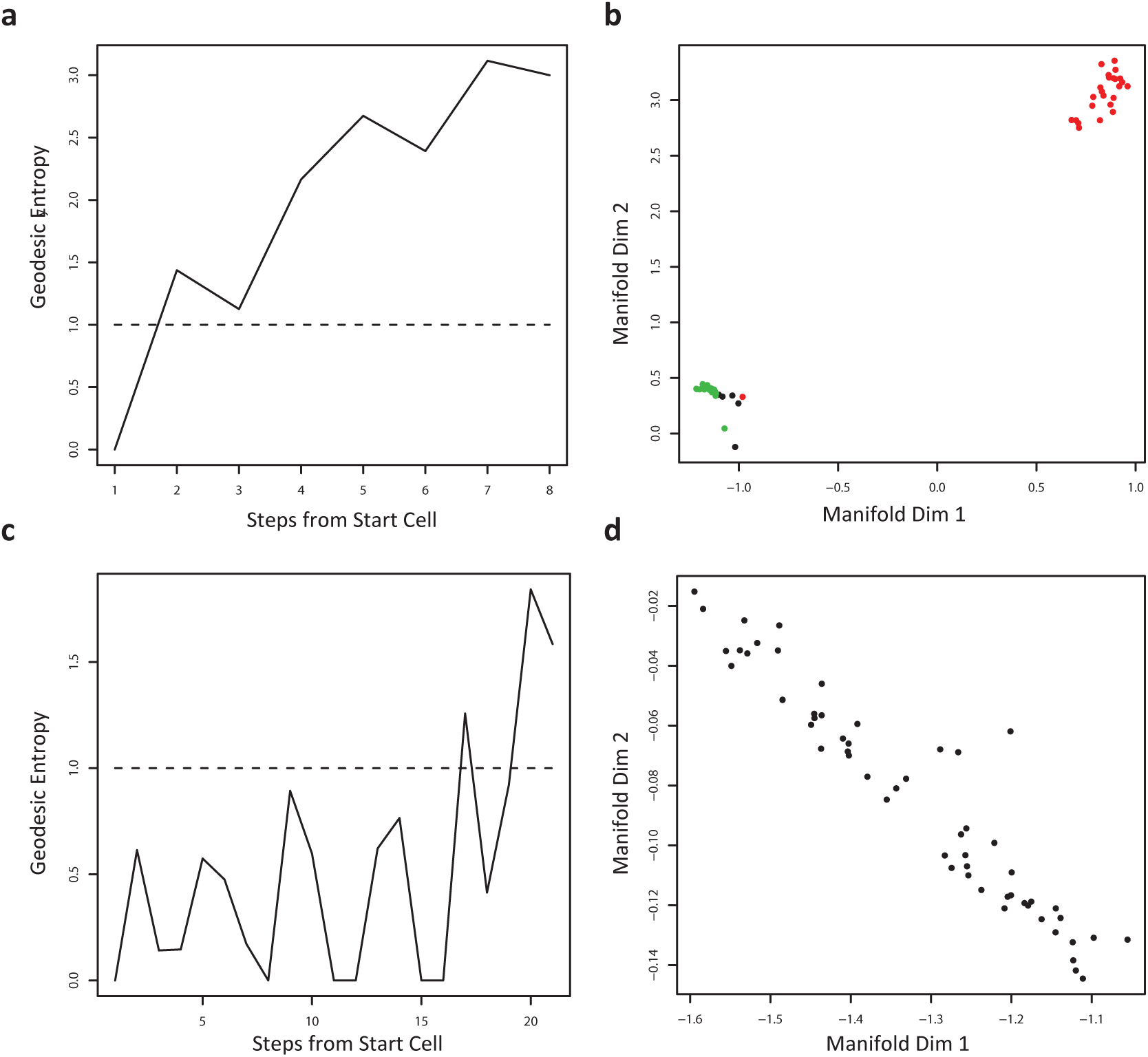
Detecting multiple branches in the mouse neural stem cell dataset. (a) Geodesic entropy computed recursively for the main trajectory branch containing neuroblasts and oligodendrocytes. Entropy exceeds 1 almost immediately indicating the presence of a second branch separating neuroblasts and oligodendrocytes. (b) Neuroblasts and oligodendrocyte cells colored by SLICER’s branch assignments. (c) Geodesic entropy computed recursively for the main trajectory branch containing active neural stem cells. Note that geodesic entropy exceeds 1 only near the end of the branch due to the small number of cells at that distance from the starting cell. SLICER does not detect a branch because the number of cells on falls below a user-specified threshold (10 by default). (d) Active neural stem cells colored by SLICER’s branch assignments.

**Supplementary Figure 8:**
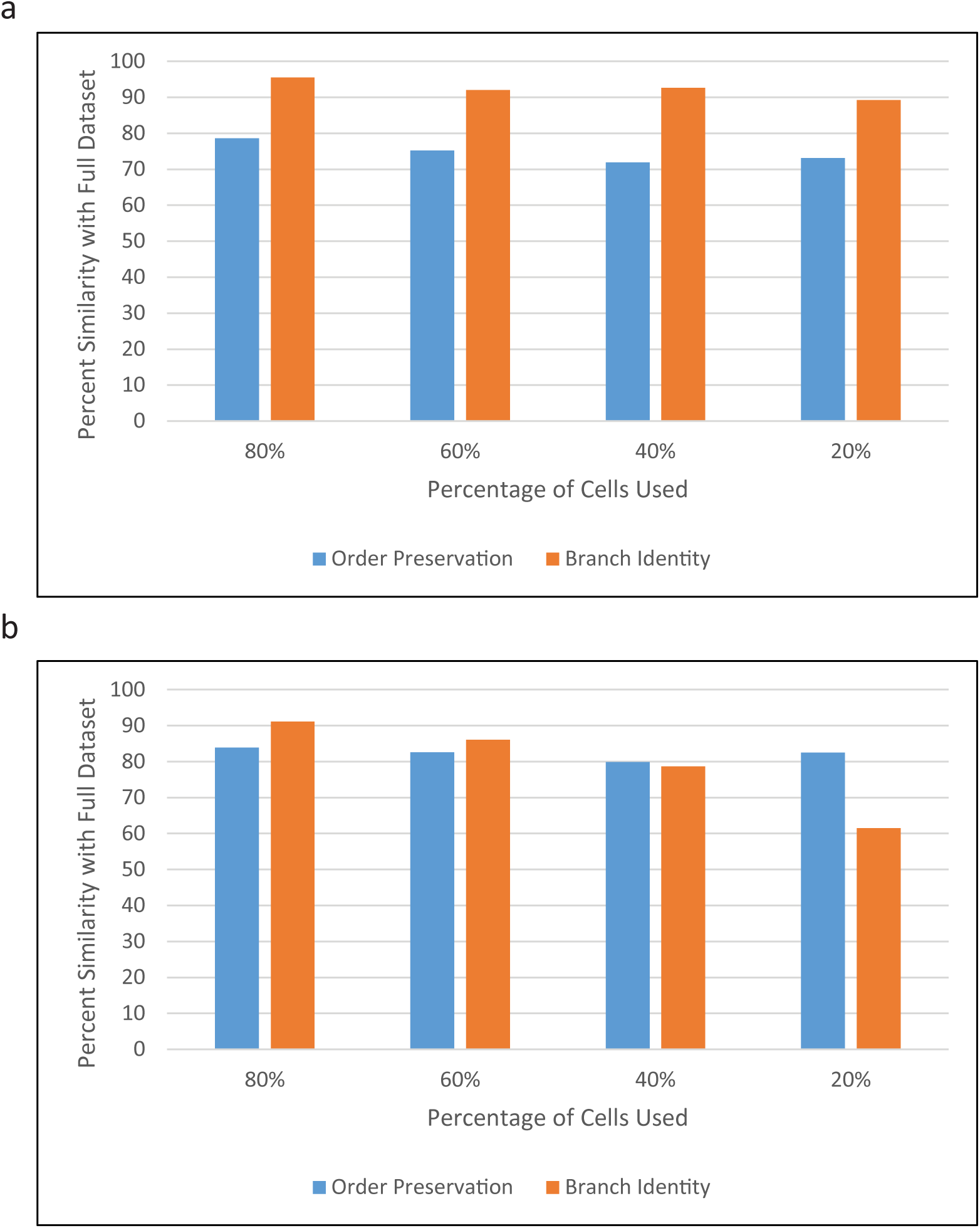
Accuracy of trajectory reconstruction using a subset of cells. (a) Graph showing how similar the SLICER trajectory is when computed using a random subset of lung cells. The blue bars show the similarity in cell ordering (units are percent sorted with respect to the trajectory constructed from all cells). The orange bars show the similarity in branch assignments (percentage of cells assigned to the same branch as the trajectory constructed from all cells). The values shown were obtained by averaging the results from 5 subsampled datasets for each percentage (80%, 60%, 40%, and 20%). (b) Order preservation and branch identity values computed as in panel (a), but for datasets sampled from the neural stem cell dataset.

**Supplementary Figure 9:**
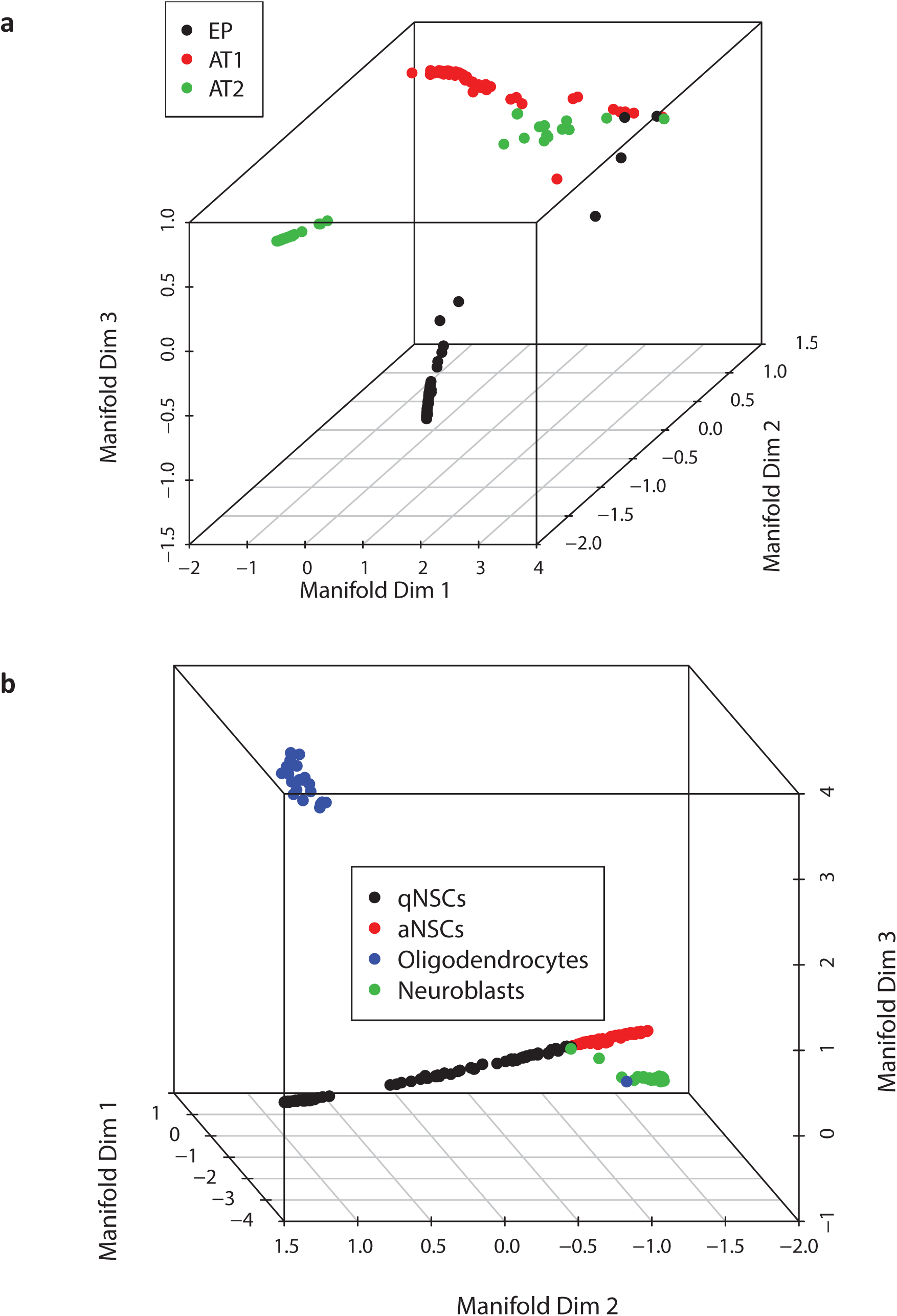
Three-dimensional HE results for biological datasets. Points are colored based on SLICER branch assignments using two-dimensional LLE embedding. (a) LLE embedding of distal lung epithelium data. (b) LLE embedding of neural stem cell data.

**Supplementary Figure 10:**
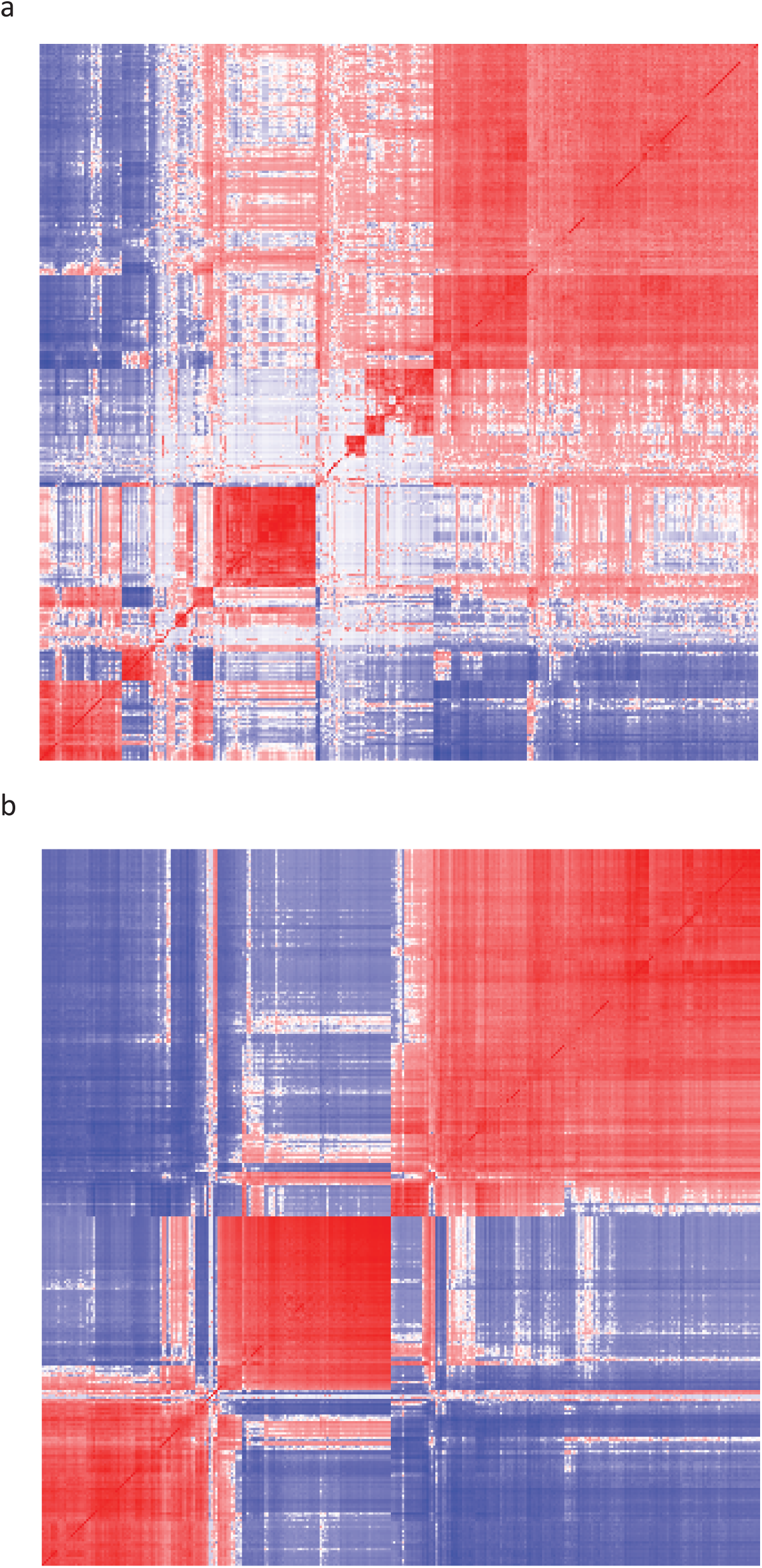
Correlation matrices for genes selected by SLICER. Blue indicates negative correlation and red indicates positive correlation. (a) Genes selected from the distal lung epithelium dataset. (b) Genes selected from the neural stem cell dataset.

## Funding

National Institutes of Health (NIH) [HG06272] to JFP, NSF Graduate Research fellowship [DGE-1144081], NIH BD2K Fellowship [T32 CA201159] to JDW.

